# Benchmarking unsupervised methods for inferring TCR specificity

**DOI:** 10.1101/2024.10.26.620398

**Authors:** Charline Jouannet, Hélène Vantomme, David Klatzmann, Encarnita Mariotti-Ferrandiz

## Abstract

Identifying T cell receptor (TCR) specificity is crucial for advancing the understanding of adaptive immunity. Despite the development of computational methods to infer TCR specificity, their performance has not been thoroughly compared. We addressed this by curating a unified database of 190,670 human TCRs with known specificities for 2,313 epitopes across 121 organisms, combining data from IEDB, McPAS-TCR, and VDJdb. Nine methods for clustering TCRs based on similarity were benchmarked against this dataset. DeepTCR demonstrated the best retention, while ClusTCR, TCRMatch, and GLIPH2 excelled in cluster purity but had lower retention. DeepTCR, Levenshtein distance, and TCRdist3 generated large clusters, with DeepTCR showing high purity. Smaller, antigen-specific clusters were produced by Hamming distance, GIANA, and iSMART. GLIPH2 and DeepTCR were the most sensitive in capturing antigen-specific TCRs. This study offers a unified TCR database and a benchmark of specificity inference methods, guiding researchers in selecting appropriate tools.

## Introduction

T-cells are characterized by the expression on their surface of a unique antigen-specific receptor, called the T-cell Receptor (TCR). The TCR is a heterodimer formed with two independently generated immunoglobulin-like chains through a recombinatorial mechanism between a collection of Variable (V), Diversity (D) and Junction (J) genes ([1]) that together recognize antigen-derived peptides bound to major histocompatibility complex (MHC) [2]. This recombination generates a highly variable region at the junction of the assembled genes called the complementary determining region 3 (CDR3), which is in direct contact with the peptide. The T-cell specificity is defined by the reactivity of a given TCR to a given peptide, entailing physiochemical interactions between the MHC/peptide complex and the V(D)J region, which engage T-cell on its activation. It is well established that T-cells are cross-reactive through their unique TCR, therefore capable of recognizing multiple antigens [3]. Unravelling this antigen-specificity is challenging due to the requirement of multi-cellular and molecular interaction. So far, TCR antigen specificity has been characterized based on preconceived in-vitro or in-vivo assays with well-established peptides, far-beyond the universe of possible interactions [3]. As such, they are not powerful enough to generalize easy and accurate screening for TCR specificity.

With the advent of next-generation sequencing of the TCR and the concomitant development of computational tools, new approaches have been proposed to infer the TCR specificity based on sequence similarity and pattern identification. These tools are all on the ground-foundation that TCRs recognizing identical peptides exhibit conserved motifs and sequences within their CDR3 regions. Several methods and tools for TCR specificity inference have been proposed. Among the most commonly used, on the one hand the Levenshtein Distance (LD) and the Hamming Distance (HD) approaches as well as TCRMatch and TCRdist3 tools have been conceived based on sequence similarity measures ([4], [5], [6], [7], [8], [9]). On the other hand, also commonly used, clusTCR, iSMART, GIANA, GLIPH2 and DeepTCR are rather focused on motif identification and clustering ([10], [11], [12], [13], [14]).

Despite their availability, each method performance has been analysed using different metrics and datasets, thus hindering the ability to compare their relative performances. A recent study by Hudson et al compared the predictive performances of part of the abovementioned tools [15] using a partially curated version of the VDJdb containing pairs of TCR alpha (TRA) and beta (TRB) chain rearrangements annotated with their antigen-specificity. Their results showed comparable performances between the tools, regardless of the model used behind for inference, with very sophisticated tools performing as well as a simple Hamming distance, for example. Moreover, the authors found that depending on the dataset (three different annotated TCR databases), the prediction performances were different. They suggested that these discrepancies could be due to different sequence pre-processing. Another possible reason is the degree of reliability of antigen-specificity annotations in these databases.

Indeed, not all TCRs from these databases were unambiguously assessed for their antigen-specificity, which could lead to multiple false positive annotations and therefore prediction. Therefore, we propose here a refined benchmarking of the most used tools on a curated database that combines pairs of TCRs from three publicly available databases, VDJdb, IEDB and Mc-PasTCR, together with unified scoring for antigen-specificity determination reliability.

## Materials and Methods

### Dataset

The dataset used in this analysis is based on a combination of three publically available databases: Immune Epitope DataBase (IEDB)[16], VDJ database (VDJdb)[17] and the manually curated catalogue of pathology associated T-cell receptor sequences (McPAS-TCR)[18], with the data being collected as of March 2023. Within the IEDB database, accessed via iedb.org, specific parameters were set : ‘Any Epitopes’ in the Epitope field, ‘T Cell’ and ‘Positive’ outcome in the Assay field, ‘Human’ as the host, ‘Any’ in the Disease field, and ‘TCR alpha/beta’ as the TCR type. To accommodate analyses based on MHC restriction types, this database was accessed twice with distinct settings for MHC restriction: ‘MHCI’ to identify CD8+ T cell related sequences and ‘MHCII” for those associated with CD4+ T cells. For both VDJdb and McPAS-TCR, filters were applied to select only data pertaining to human species. Notably, within these two databases, every sequence is documented with information on the cell subset. Subsequently, these three databases were combined into a unique dataset, undergoing extensive curation for uniformity of information. In cases where duplicate and triplicate sequences between the three databases shared identical information (same V-CDR3 amino acid-J for both TRA and TRB, same epitope, same organism, same PubMed ID and same cell subset), only one instance was retained (486 sequences are involved). If there were missing information or discrepancies between the databases for the same source, each instance was preserved to leave the choice to the user.

For each TRA/TRB pair, a sequence reliability score, named *Verified_score*, was established. The verification process of the sequence was primarily conducted using the IEDB, relying on its comprehensive curation strategy [16]. We assessed the concordance between the ‘calculated’ and ‘curated’ sequences by considering the start and end positions of the CDR3 sequences, as well as the delineation at the ‘C - F’ border. For the other two databases, assignment of information was contingent upon the availability of both the alpha and beta CDR3 sequences. A score of 2 indicates both TRA and TRB sequences are known and verified; a score of 1.1 is attributed when only the TRA sequence is present and verified and a score of 1.2 when this is the case for the TRB sequence, finally a score of 0 is set when neither sequences are verified. Similarly, the method of antigen identification is used to assign a second score, *Antigen_identification_score*, allowing users to filter on this parameter as detailed in **Supplementary Table 1**.

For the purpose of the benchmarking, we selected only the sequences with a *Verified_score* of 2 and an *Antigen_identification_score* above 4.3. Additional filters included: (i) selection of pairings with known V and J information, (ii) exclusion of pairings with unknown epitope, (iii) selection of TCRs with CDR3s of 6 to 23 amino acids, and (iv) selection of epitopes that bind at least 2 TCRs defined as a unique CDR3a/CDR3b pair. Moreover, TCR originating from 10X genomics datasets were excluded from the combined database. The resulting dataset comprises 5 261 TCRs from CD8+T-cells and 192 TCRs from CD4+T-cells, all with known VJ and epitope information. The curation strategy of this pooled database is depicted in **Figure 1A**.

**Figure 1:**
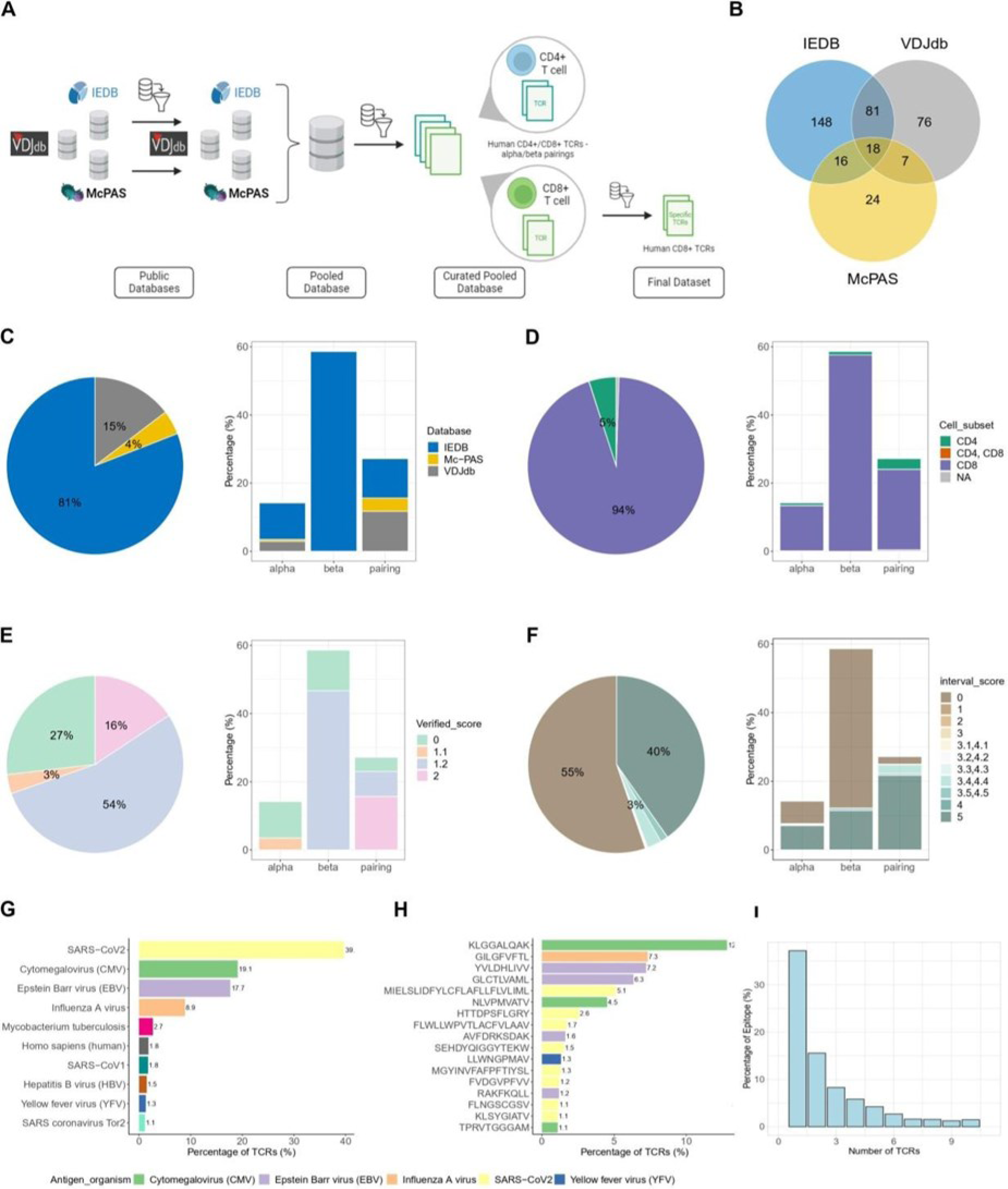
Overview of the pooled database composition. (A) Curation strategy: human TCRs extracted from three public databases, IEDB, VJDdb and Mc-PAS are combined into a large dataset. After applying several filters (see material and methods), the pooled database is filtered according the cell type CD8+T-cells. (B) Venn diagram representing overlap of studies across the three public databases. (C) Database contribution: overall distribution of public databases in the final pooled database in the whole data (left panel), and their distribution across different chain types: alpha, beta and alpha/beta pairings (right panel). (D) Cell type distribution: in the entire pooled database in the whole data (left panel) and across each chain type: alpha, beta and alpha/beta pairings (right panel). (E) Verified score distribution across the entire pooled database (left panel) and within each chain type (right panel). (F) Antigen_identification_score distribution in the entire pooled database (left panel) and within each chain type (right panel). (G) Predominant antigen organisms: barplot illustrates the most represented antigen organisms in the pooled database, with number indicating the percentage of TCRs binding to each organism. (H) Key epitopes: barplot displays the most represented epitopes in the pooled database, with number reflecting the percentage of TCRs binding to each epitope. (I) Epitope-TCR binding analysis: barplot showing the distribution of epitopes in the pooled database based on number of TCRs binding to them.

### Benchmarked tools and approaches

We benchmarked the performance and the clustering accuracy of existing methods. We focused our analysis on nine approaches and tools: Levenshtein Distance (LD) [4] [5], Hamming Distance (HD) [6], [7], TCRMatch [8], TCRdist3 (v.0.2.2) [9] GIANA (v.4.1) [12], iSMART [11] clusTCR (v.1.0.2) [10] GLIPH2 [13] and DeepTCR (v.2.0) [14]. Note that only GLIPH2, DeepTCR, TCRdist3 and ClusTCR can integrate paired TRA and TRB as input. Hence, for LD, HD, TCRMatch, GIANA and iSMART, TRA and TRB rearrangements were analysed independently, whereas for TCRdist3, clusTCR, GLIPH2 and DeepTCR both TRA and TRB rearrangements from paired TCRs were considered. Additional information regarding each approache and the parameters used are detailed in Supplemental Info.

### Clustering performance and similarity metrics

Assuming that the similarity of CDR3 sequences correlates with epitope specificity, some metrics were set out to assess the effectiveness of the clustering approaches mentioned above. These metrics include ‘purity’, ‘retention’ and sensitivity, which evaluate the quality of the clusters formed. In details:

**Retention** is defined as the ratio of the number of clustered CDR3 sequences to the total number of sequences in the dataset. This measure represents the fraction of CDR3 sequences successfully clustered by the applied method.

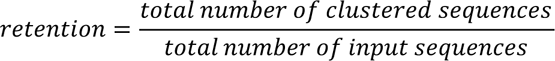

**Purity** is defined as the fraction of CDR3 sequences within a single cluster targeting the same epitope. Purity is calculated as the sum of CDR3 specific for the most represented epitope within each cluster divided by the total number of sequences in any cluster. The purity values range from 0 to 1, where 0 indicates non-pure clusters (all CDR3s bind to different epitopes) and 1 pure clusters (all CDR3s bind to a same epitope).

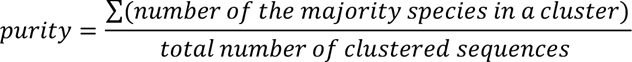

Based on the purity score, we further calculated:

- the percentage of clusters with a purity over 0.9, defined as the proportion of clusters in which the predominant epitope is the most represented (more than 90% of the cluster). For each cluster, individual purity is calculated as

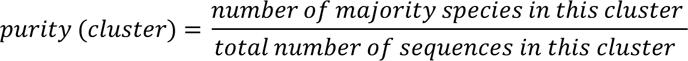

Then, the percentage of clusters with a purity >0.9 is determined as follows:

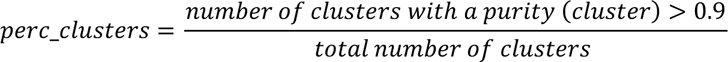

- the percentage of CDR3s in clusters with a purity over 0.9, defined as the fraction of sequences that belong to the clusters previously identified as having a purity greater than 0.9.

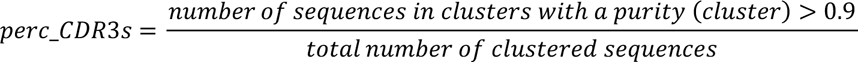

**Sensitivity** is defined as, for a given epitope, the fraction of epitope-specific CDR3s contained within epitope-specific clusters. Epitope-specific clusters are selected based on two criteria: a size greater than 3 sequences and a number of epitope-specific sequences at least twice higher than the other sequences in the cluster. Then, the sensitivity for each epitope is calculated as:

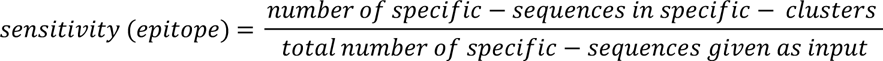

Retention, purity, including the percentage of clusters with a purity greater than 0.9 and the percentage of sequences/pairs belonging to these high-purity clusters (as previously described) as well as sensitivity were computed on the output of each method. First, these metrics were calculated on the global clustering outputs on the pre-filtered curated database (filters mentioned above). Then, the analysis was extended to epitope-specific sequences. For this purpose, we selected six epitopes as detailed in the result section, each represented at various extent in the input database.

### Statistical analysis

All statistics analyses were performed using R software version 4.1. Comparative figures were produced using ggplot2 R package [19], networks were produced using Cytoscape software version 3.10.1 [20] and explicative schemas were created with <u>biorender.com</u>.

## Results

### Public TCRs are enriched for CD8 T cell TCRs and viral antigen specificities

To assess the performance of antigen-specificity inference methods of αβ TCRs, we combined and curated three well-established public databases: IEDB ([16]), McPAS-TCR ([18]) and VDJdb ([17], [21] (**Figure 1A**). These three databases were composed of TCRs, i.e. of unique CDR3a/CDR3b pairs, from overlapping studies (**Figure 1B**) representing nearly 25% of the total number of TCRs (for the remaining 75%, the source is not indicated). Specifically, 18 studies were common across the 3 databases, 81 shared between IEDB and VDJdb, 16 between IEDB and McPAS-TCR and 7 between VDJdb and McPAS-TCR.

IEDB contributes to 81% of the pooled database, followed by VDJdb at 15%, and McPAS at 4% (**Figure 1C**, left). In detail, VDJdb and McPAS-TCR provide predominantly αβ TCR pairs (11.7% and 3.9%, respectively), some α-only chains (2.9% and 0.6%, respectively), but no β−only chains (**Figure 1C)** while IEDB is mainly composed with β-only chains. In addition, 94% of the publicly available TCRs originates from CD8+T-cells (**Figure 1D**). An overview of the most representative antigen organisms and epitopes for which the TCRs were annotated (greater than 1%) and included in the pooled database is presented in **Figure 1G and H**. The predominant organism is SARS-Cov2 (approximatively 40%), followed by the Cytomegalovirus (CMV) (19%) and Epstein-Barr virus (EBV)(18%) (**Figure 1G**). The top three epitopes are KLGGALQAK (CMV), bound by 12.8% of TCRs, GILGFVFTL (Influenza), and bound by 7.3%, YVLDHLIVV (EBV), bound by 7.2% of the TCRs present in the curated pooled database. **Figure 1I** illustrates that most epitopes (>35%) are associated with only one TCR. These epitopes and their associated TCRs were excluded from further clustering analyses. Altogether, we assembled a pooled and unified database of 190,670 TCRs among which 184,137 TCRs recognize 2,313 known epitopes from 121 organisms.

### Unified scoring reveals database heterogeneity and provides tool for informed TCR selection

The TCRs populating these databases were originally identified with different degrees of sequence reliability (verification of the original nucleotide and amino-acid sequences for each TRA and TRB rearrangement) and antigen-specificity validation (e.g. cell culture with peptide or full protein, tetramer/dextramer cell sorting, expansion in or association with disease). To overcome the heterogenous level of sequence and antigen-specificityreliability, we extended the scoring on sequence validation provided by IEDB (assessed by the Verified_Score (VS)) and developed a complementary scoring method that accounts for technical approach used to assign antigen-specificity (assessed by the Antigen_identification score (AIS)) to our pooled database (detailed in Dataset). 25% of the TCR entries were classified as unreliable (VS = 0, meaning that the sequences were not double-checked). Moreover, there are 54% of reliable TRB sequences (VS = 1.2), much higher than the 3.5% of validated TRA sequences (VS = 1.1). Finally, while only 15% of the TRA/TRB pairs are unreliable and 20% of TRB-only chains, the proportion of unreliable sequences is higher for α-only chains (75%). Nevertheless, some TRA/TRB pairs feature unreliable TRA (VS = 1.2, 27%) (**Figure 1E**). Regarding antigen-specificity validation, 55% had AIS of 0, meaning that no information was available regarding the method used to assess the antigen-specificity. Conversely, 40% of the sequences have an Antigen_identification_score of 5 (**Figure 1F**, left panel). Focusing specifically on TRA sequences, we observed roughly equal proportions of sequences with no (48.1%) and high AIS (51.9%). In contrast, nearly 80% of TRB chain sequences are categorized as unreliable. Conversely, for the TRA/TRB pairings, the majority of sequences (about 80%) exhibits high AIS. Altogether, our unified scoring revealed the heterogeneity of the quality of the information available for the sequences stored in the databases and now offers an objective process to select the TCRs of interest and evaluate the performance of antigen-specificity inference tools.

### Most of the antigen-specificity inference methods form small clusters

Given the bias toward TCRs originating from CD8 T-cells in the unified database, our benchmarking analysis was focused on those predominant TCRs, further narrowed down to only those with aVS=2 and an AIS>4.3. As such, the actual analysed dataset was composed of 4,779 unique TRA/TRB pairs, including 4,103 unique CDR3a and 4,292 unique CDR3b (8,395 unique sequences). For the LD, HD, TCRMatch, iSMART and GIANA methods, we analysed only the CDR3 sequences, as per the method original design, and considered the two chains separately. In contrast, for the GLIPH2, clusTCR, DeepTCR and TCRdist3 methods, we inputted the paired CDR3a/CDR3b sequences along with the required additional information in line with their original description (see details in **Supplementary Table 2**).

By assessing the number of clusters formed and their size (**Figure 2A**), we found that ClusTCR creates the least number of clusters (111) while DeepTCR creates the highest number (1628). The size distribution of the cluster is comparable across all the methods, with the majority (>70%) of the clusters being small (< 4 sequences), suggesting a low similarity among sequences in the database. The size of the largest cluster varies from 14 (TCRMatch) to more than 150 (LD) CDR3/pairs, with TCRdist3, GIANA, iSMART and clusTCR generating cluster no larger than around 40 CDR3/pairs. Interestingly, these differences can be observed despite a similar number of clusters (e.g. LD vs. TCRdist3, GIANA and iSMART), highlighting differences in the clustering parameters. When we ran the clustering on the full database with mixed CDR3a and CDR3b, we found that LD, HD,TCRMatch, iSMART and GIANA, originally not designed for TRA/TRB pairing input, formed pure clusters composed of either CDR3a or CDR3b, as illustrated by the network visualization (**Figure 2B**, **C**). Networks visualization for clusTCR, GLIPH2, DeepTCR and TCRdist3 are also available in **Supplementary Figure 1**.

**Figure 2:**
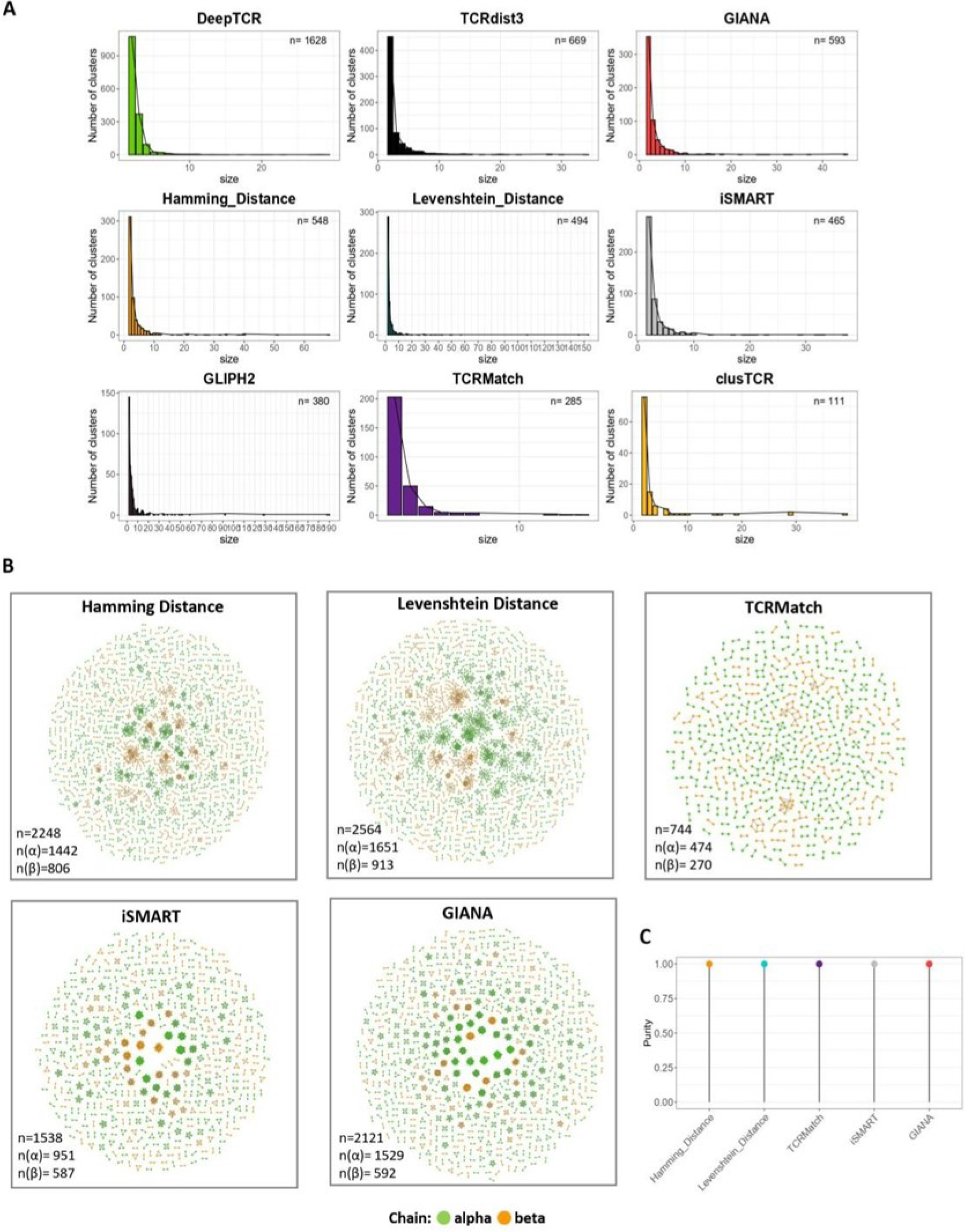
Comparative clustering analysis of the nine methods. (A) Cluster size distribution for the nine methods, ordered by decreasing number of clusters (B) Network visualization of output clusters: this section displays networks generated by five methods - Hamming distance (HD), Levenshtein distance (LD), TCRMatch, iSMART and GIANA. Each subnetwork represents a cluster, with individual dots symbolizing CDR3 sequences; green denotes CDR3a and orange the CDR3b. The connectivity criteria vary for each method: HD=1 for HD (top-left panel), LD=1 for LD (top-middle panel), a TCRMatch score higher than 0.97 (top-right panel) and manual linkage of sequences within the same cluster for both iSMART (bottom-left panel) and GIANA (bottom-middle panel). (C) Purity analysis: this graph illustrates the purity of clusters generated by the five first methods according to the chain type.

### DeepTCR shows the highest retention and clusTCR and GLIPH2 the highest purity for TCR clustering

To compare how antigen-specificity is captured by the clustering approaches, we computed theretention, as the number of sequences actually clustered out of the total number of input sequences, andthe purity of the clusters, as the fraction of the sequences annotated with the most abundance antigen specificity covered among all clusters (see Materials and Methods and **Supplementary Figure 2**). **Figure 3A** summarizes the results for each metric across all methods and **Supplementary Table 2** details the number of sequences clustered. LD, HD, GIANA, and iSMART, showed similar low retention (0.25 +/- 0.06) and intermediate purity scores (0.66 +/- 0.09) whereas TCRMatch displayed a very low retention (0.09) and much higher purity (0.82). DeepTCR clusters outperform all the methods in terms of retention (0.92) and displayed an intermediate purity (0.63). clusTCR clusters the lowest number of sequences (retention = 0.09) with a purity of 0.99, similarwith TCRMatch. GLIPH2 ranks in the third place regarding the purity (0.8) with a low retention to 0.22.Surprisingly, TCRdist3 stands out for balancing both metrics (retention = 0.44, purity = 0.42). In summary, we found a negative modest correlation between purity and retention (**Figure 3B**), revealing subtleties driven by each method. These results suggest that TCR with close enough sequences can recognize distinct epitopes, as supported by recent studies ([22], [23], [24]).

**Figure 3:**
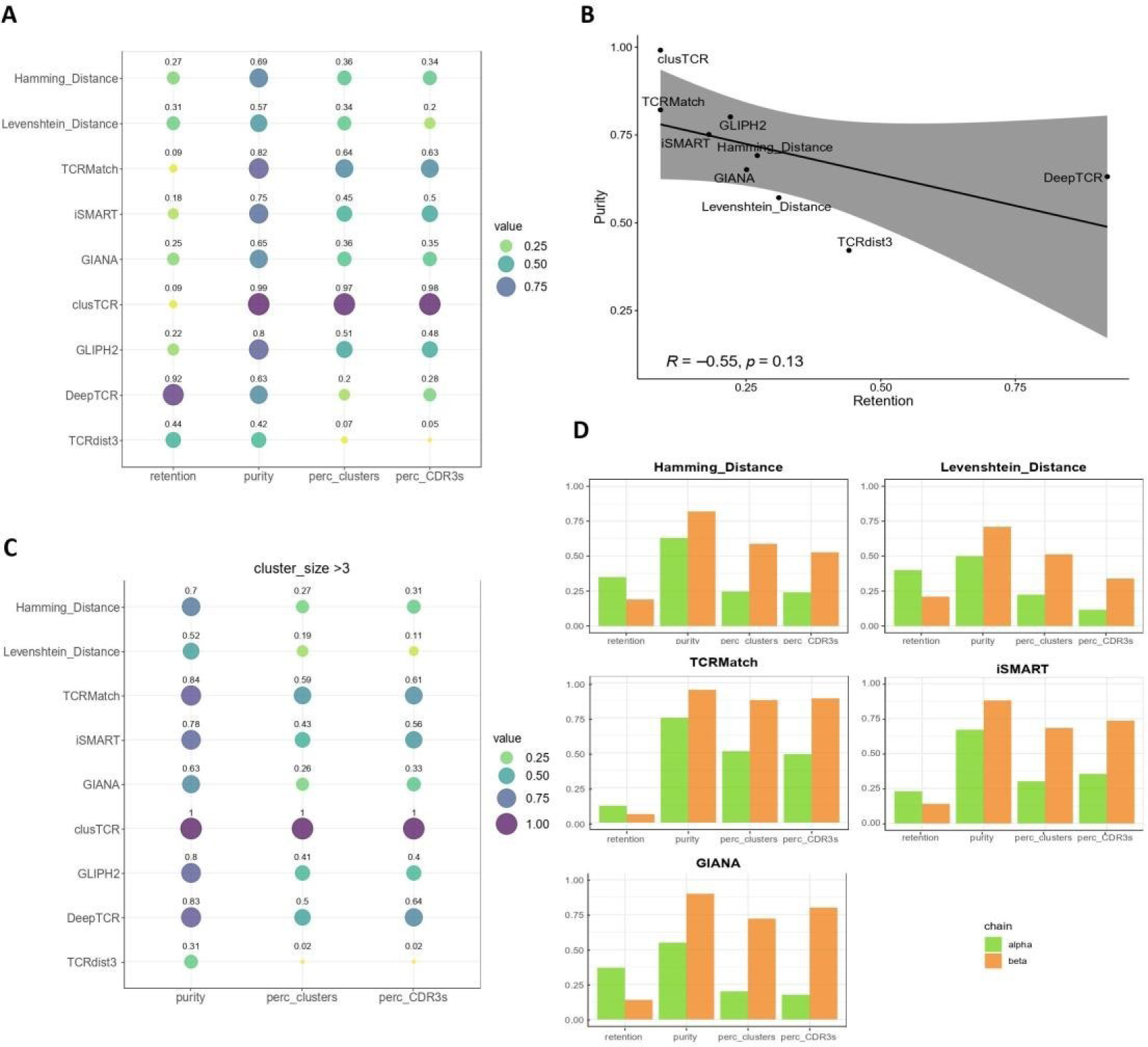
Evaluation of method performance. (A) Performance metrics: bubble plot showing different performance metrics for the nine methods, including retention, the purity, the percentage of clusters with a purity over 0.9 and the percentage of sequences within these clusters. All metrics are normalized between 0 and 1 (with the last two metrics needing multiplication by 100 to convert their values into percentages). (C) Correlation plot of the purity as a function of the retention. P-value was calculated with a Pearson test. (B) Same analysis as in (A) but with a focus on clusters with more than 3 sequences. (C) Specific metrics: retention, purity, percentage of clusters with a purity over 0.9 and percentage of sequences within these clusters for the HD, the LD, TCRMatch, iSMART and GIANA, analysed separately for alpha and beta clusters

We further quantified the fraction of highly pure clusters (at least 90% of purity) and the sequences/pairs forming these clusters. Interestingly, 97% of ClusTCR clusters are of at least 90% purity, capturing 98% of the sequences clustered. TCRMatch, GLIPH2 and iSMART showed intermediate scores (64%, 51% and 45% clusters with at least 90% purity, respectively; and capturing 63%, 48% and 50% of the sequences, respectively), followed by LD, HD and DeepTCR (34%, 36% and 20% of highly pure clusters capturing 34, 20% and 28% of the sequences, respectively). Conversely, TCRdist3 has the lowest score values (7% of 90% purity clusters; 5% of the pair). Of note, when the TRA and TRB chains were analysed separately using HD, LD, TCRMatch, iSMART and GIANA, we found that both the retention and the purity are lower for the TRA chain than the TRB chain (**Figure 3A**). Given that all the sequences were initially from reliably annotated pairs of TCRs, this result suggest a differential contribution of each chain to antigen specificity, as proposed earlier ([25]).

However, as shown in Figure2A, most of the methods generate predominantly small clusters made of 2 to 3 sequences, a bias that can influence the global purity score. We then computed the purity considering clusters composed by strictly more than 3 sequences or pairs (**Figure 3C).** Purity remained consistent with previous results for most of the methods, except for LD, TCRdist3 and DeepTCR (0.52, 0.83 and 0.31 respectively). For the LD and TCRdist3, while overall purity was comparable, we found a two-fold decrease in high purity clusters percentage (19% and 2% respectively) and in the percentage of sequences in these clusters (11% and 2%), revealing that the purest clusters were those formed by at most 3 sequences, while larger clusters are formed with sequences with different specificities. For DeepTCR, the overall purity (0.83), the percentages of high purity clusters (50%) and of pairs in these clusters (64%) increased, suggesting that clusters smaller than 3 sequences mainly were heterologous. These results were further confirmed when considering clusters larger than 5 and 10 sequences (**Supplementary Figure 3**). The number of clusters formed when considering clusters size >3, 5 or 10 sequences/pairs is shown in **Supplementary Table 3**.

Altogether, our results indicate major differences regarding sequence/pair retention depending on the methods, with DeepTCR being the most conservative, TCRMatch the least.

### GLIPH2 and DeepTCR show highest sensitivity in epitope-specific TCR clustering

We further analyzed the sensitivity of each method selected for the GIL-epitope focusing on clusters where the GIL-sequences/pairs predominated (**Figure 4A**, **Supplementary Figure 6)** as detailed in the Materials and Methods section (**Figure 4B**). GLIPH2 followed by DeepTCR exhibited the highest sensitivity, indicating their effectiveness in capturing GIL-specific pairs. Conversely, TCRdist3 showed the least sensitivity. When comparing the sensitivity across the six selected epitopes, GLIPH2 demonstrated the highest median sensitivity (∼0.4), closely followed by DeepTCR (∼0.35), whereas TCRdist3 and TCRMatch have the lowest value (**Figure 4C**). Overall, GLIPH2 and DeepTCR emerged as the most effective method for capturing epitope-specific sequences.

**Figure 4:**
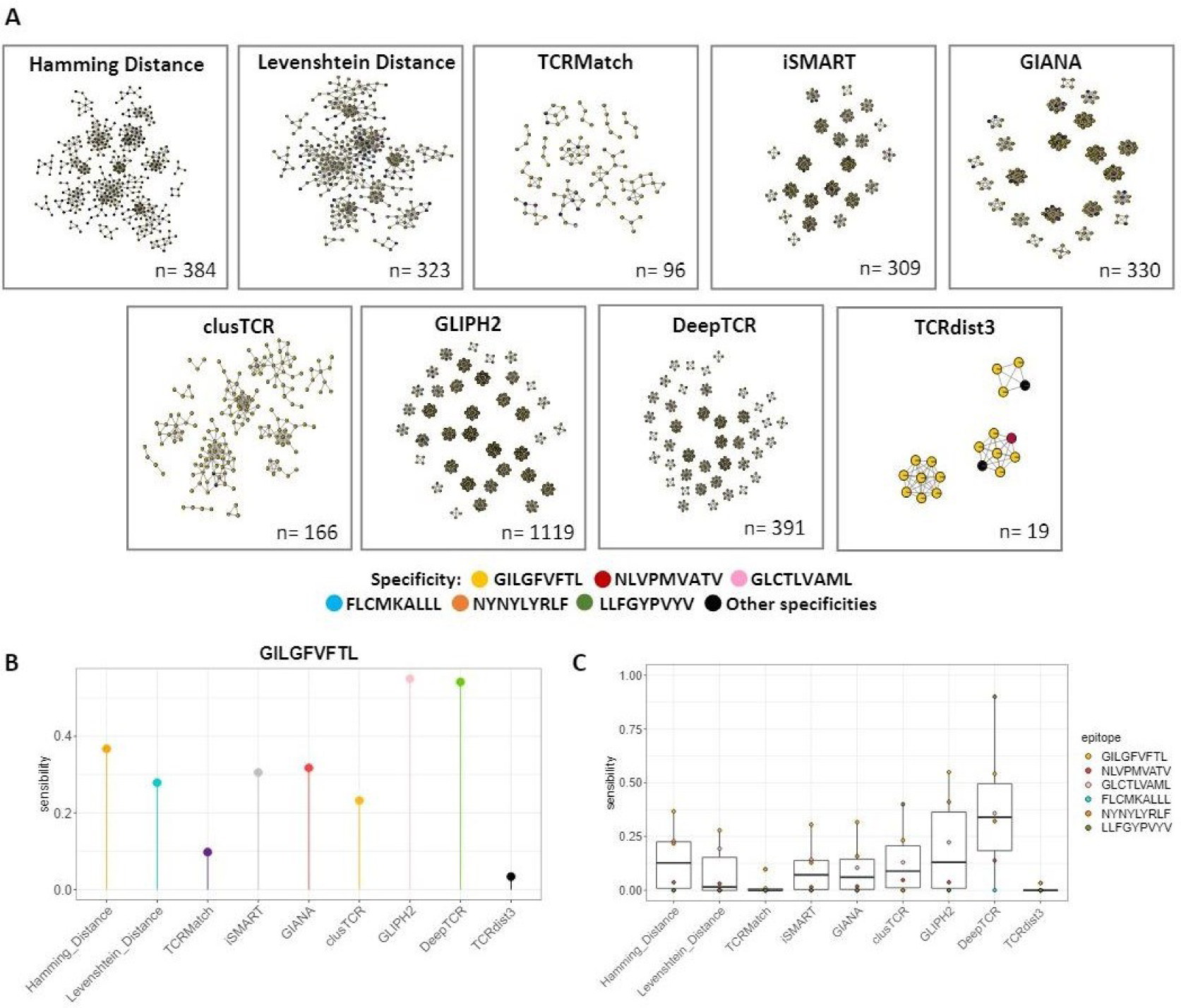
Focused analysis on GILGFVFTL-specific clusters and sensitivity. (A) Network visualization of the purest clusters representing clusters selected where GILGFVFTL is the predominant epitope, being at least twice as abundant as the second most used epitope in the cluster across each method. (B) Sensitivity of the GILGFVFTL epitope for each method. (C) Epitope wide-Sensitivity comparison for the 6 selected epitopes across each method. A Wilcoxon-test was released but there is no significance.

### Poly-specific TCRs are captured by most of the methods

We further quantified the numbers of specificities per cluster. More than 70% of the clusters formed by HD, LD, GIANA, iSMART, DeepTCR and TCRdist3 assemble TCRs with at least 2 specificities, i.e. in polyspecific clusters, whereas TCRMatch, clusTCR and GLIPH2 predominantly formed monospecific clusters (**Supplementary Figure 4**). The distribution of the antigen-specificities in the database is far from being even. As such, some specificities are highly represented, such as GILGFVFTL (later named GIL) associated with 706 distinct TRA/TRB pairs, as compared to others relatively less represented, like LLFGYPVYV associated with only 10 pairs (**Supplementary Table 4** and **Figure 5A**). To evaluate the impact of such imbalance, we focused our analysis on the clusters formed by each method with the most represented epitope (GIL). Consistent with the overall retention, DeepTCR clusters nearly 100% of the GIL-specific annotated pairs while TCRMatch followed by ClusTCR grouped less than 30% of the same pairs (**Figure 5B**). HD, LD, GIANA and iSMART retains about 50% of the sequences. GLIPH2, despite a moderate retention score, captures approximately 67% of GIL-specific pairs. Importantly, for all the methods, half or more of the GIL-specific clusters are formed by at most 3 sequences (**Figure 5C**).

**Figure 5:**
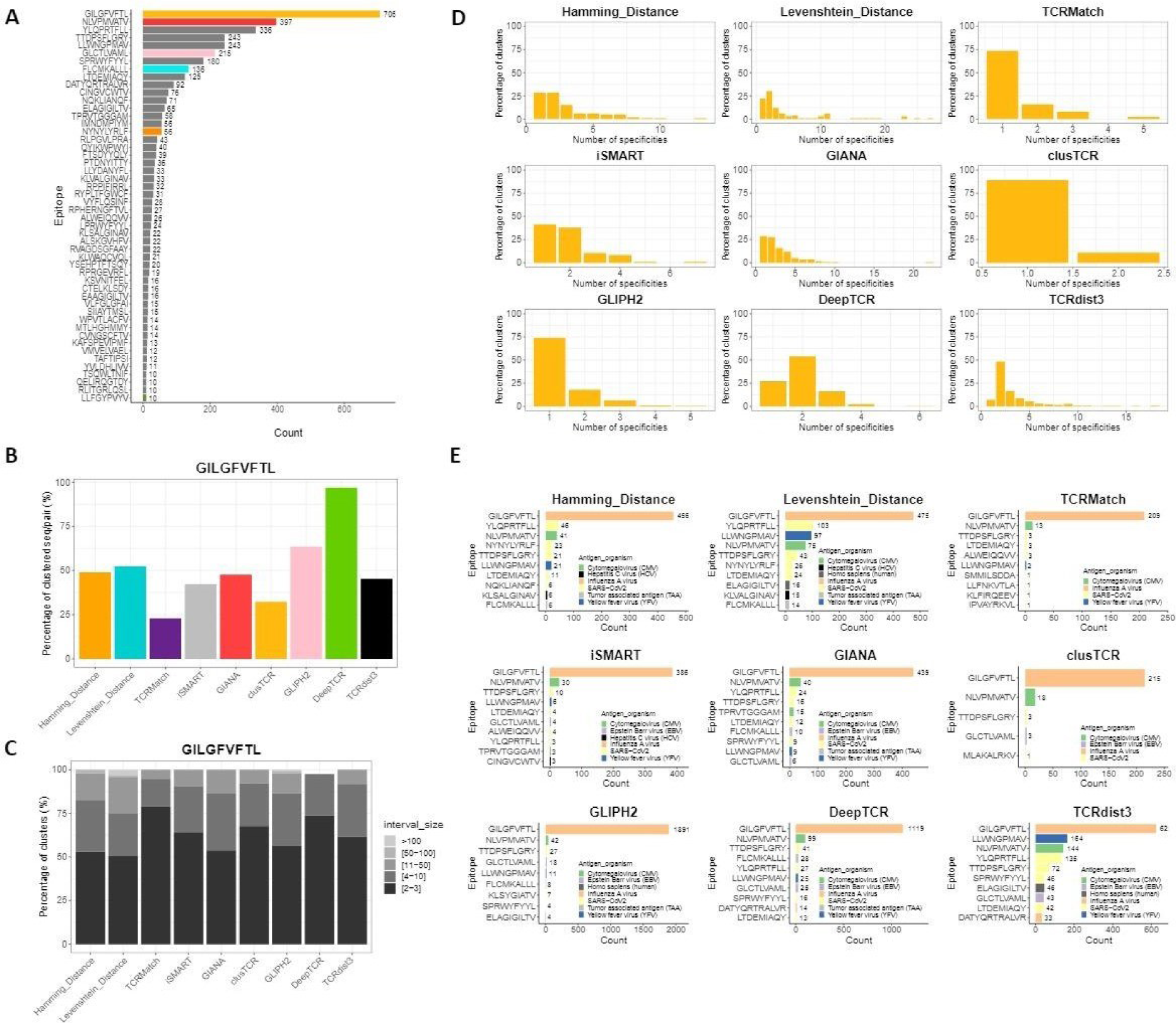
Analysis of method performance using specific epitopes. (A) Representation of top epitopes, most represented epitopes in the input data, focusing only on those with a frequency greater than 1%. Six epitopes were selected based on their frequency (both high and low) and are color-coded as follow: dark gold for GILGFVFTL, red for NLVPMVATV, pink for GLCTLVAML, blue for FLCMKALLL, orange for NYNYLYRLF and olive for LLFGYPVYV. (B) Percentage of GILGFVFTL-specific-sequences/pairs clustered by each method. (C) Size interval distribution in GILGFVFTL-specific clusters for each method. (D) Specificity distribution in GILGFVFTL-specific clusters as a function of the number of specificities contained in each cluster, for each method. (E) Top additional epitopes in GILGFVFTL-specific clusters displaying the top 10 (or fewer) additional epitopes present in these clusters for each clustering method, coloured by antigen organism. The numbers displayed on the barplot correspond to the total number of sequences specific to the different epitopes associated with the specific GILGFVFTL clusters.

We then further analysed the distribution of the other specificities present in the GIL-labeled clusters. TCRMatch, clusTCR and GLIPH2 predominantly formed monospecific GIL-specific clusters whereas HD, LD, iSMART, GIANA, DeepTCR and TCRdist3, favored polyspecific clusters (**Figure 5D**). The distribution of the other specificities present in the GIL-clusters was strikingly comparable between all the methods generating polyspecific clusters (**Figure 5E**), with the epitopes most frequently identified among all nine methods being YLQPRTFLL, TTDPSFLGRY (from SARS-Cov2), NLVPMVATV (from CMV) and LLWNGPMAV (from Yellow Fever virus). The same trends were found when analyzing clusters labelled with the least represented epitope in the database (FLCMKALLL, **Supplementary Figure 5**). These observations suggest that the ability of the method to capture antigen-specific sequences, including clustering poly-specific TCRs, is independent on the abundance of the antigen-specific TCRs in the database.

## Discussion

Given the exponential increase in the generation of TCR sequences, it would be of utmost importance to be able to predict their antigen specificity. Following significant progress in the field of bioinformatics, many tools have been developed by various teams to infer TCR specificity using different strategies such as theoretical frameworks, computational algorithms, and deep learning approaches. In this study, we conducted an independent comparison of the predictive capabilities of some unsupervised sequence-based methods. Most of them rely on assessing global and/or local similarity, while others harness deep learning process to extract meaningful features from AA sequences. We explored the efficacy of selected clustering methods in categorizing αβTCR proteins according to their specificity, namely their ability to recognize epitopes. Notably, all methods benchmarked in this study either exclusively focus on or attribute significant importance to the CDR3 region, highlighted by its critical function in peptide binding.

This analysis required input data where the assignment of specificity to TCR sequences was reliable. To this end, we established a harmonized database from three public databases that, while enriched manually by authors of articles addressing this topic, are prone to errors ([16], [18], [21]). This consolidated database empowers users to choose sequences by evaluating both their inherent accuracy and the dependability of their specificity designation, thereby guaranteeing data of superior quality for analysis.

Using this unified database, we observed that such as LD, HD, TCRMatch, GIANA and iSMART, not integrating TRA/TRB pairs as well as clusTCR and GLIPH2 partially integrating TRA/TRB pairs, achieved low retention and high purity as compared with methods integrating TRA/TRB pairs. Regarding antigen-specificity, methods based on distance measurements with a threshold of one, such as LD or HD, or relying on sequence alignment, including iSMART or GIANA, tended to form both poly-specific and mono-specific clusters. In contrast, approaches based on k-mers such as TCRMatch or motifs such as clusTCR and GLIPH2 tend to generate mono-specific clusters. In contrast, TCRdist3 achieved suboptimal performance and predominantly forms poly-specific clusters. A similar trend was observed with deep learning methods, such as DeepTCR, which also favored the formation of poly-specific clusters but achieves better performance in terms of purity than TCRdist3. It is established that a single TCR may recognize multiple distinct epitopes [22], [26]. Example with the pair CATDTTSGTYKYIF (CDR3 alpha) CSARDLTSGANNEQFF (CDR3 beta) which recognizes multiple peptides in the pooled human database, as ENPVVHFFKNIVTP (maltose binding protein, Homo sapiens), IIPAFHFLKSEKGL (androglobine, Homo sapiens) or DVSKVHFFKGNGQT (transporter atp-binding protein, Rhizobium leguminosarum) [27]. Furthermore, the specificity of a TCR can vary significantly when a TRB is paired with different TRA [22]. Moreover, TCRs with completely dissimilar amino acid sequences may bind to the same epitopes. For example in the pooled database, the CASSLLGGWSEAFF (CDR3b) - CAASHIQGAQKLVF (CDR3a) TCR and the CASSIRSSYEQYF(CDR3b) – CAAGGSQGNLIF (CDR3a) TCR bind both the GILGFVFTL epitope [28]. To be more specific, 12 954 distinct TCRs are known to bind the GILGFVFTL epitope. Importantly, the mere binding of a TCR to an epitope does not guarantee T-cell activation [29]. Given that poly-specificity is a natural and very common phenomena ([3]), having access to larger datasets of TCR pairs experimentally validated for their poly-specificity should help improve the methods and models behind.

Our analysis revealed a noticeable difference in metric outcomes when methods do not utilize paired TRA/TRB chain. Specifically, TRA chain sequences are more often organized into clusters than their TRB counterparts. Furthermore, the TRA chain clusters typically tended to be more polyspecific compared to those formed from TRB sequences. A previous study of the VDJdb identified overrepresentation of dual TRA expressing cells, determining whether the highly clustering TCRs belongs to that category would shed light on this particular phenomenom and the associated immune response [30].

This point is particularly important given that some methods in our study cluster TCRs by considering both the TRA and TRB chains, while others integrate only beta chains. Notably, even though tools like ClusTCR, TCRdist3, and GLIPH2 have theoretically the capability to process data from paired TCRs, they are not optimized to perform paired chain clustering. Indeed, GLIPH2 assigns clusters based on the CDR3b region only, clusTCR generates an encoding for TRA and TRB sequences individually and then merges them and TCRdist3 calculates a distance matrix for each chain and then adds them together. To determine the effectiveness of incorporating both TRA and TRB chain information, a detailed examination across these models, including DeepTCR, is necessary. This could highlight the unique strengths of each method, particularly valuable in the analysis of complex single-cell sequencing data. When evaluating the performance of each method for six epitopes selected from different species and expressed differently in the filtered pooled database, we observed that TCRMatch and TCRdist3 demonstrated variability in their clustering outcomes forming both monospecific and polyspecific. Nevertheless, this selection enabled us a valuable perspective on how each method adapts to the epitope’s representation within the dataset. Furthermore, our findings indicate that GLIPH2 and DeepTCR exhibit superior sensitivity across the six epitopes, adeptly identifying sequences sharing the same specificity.

The methods studied in this manuscript are sequence-based. As such, the representation of AA sequences, while potentially increased with physicochemical properties and/or biological features, does not fully encapsulate the entirety of relevant information. Structure-based specificity inference models considering the three-dimensional architecture of the TCR-peptide/MHC complex would be instrumental for determining both binding between the two entities [31]. Recent studies posited that hybrid models combining sequence and structure-based methods could enhance performance and yield more accurate specificity predictions [32]. Nevertheless, the lack of three-dimensional structural data in public repositories presents a significant obstacle. To address this gap, tools like TCRmodel2 have emerged, using AlphaFold [33], [34] to predict structures directly from AA sequences, thus offering a practical solution to overcome this challenge [35]. Interestingly, other methods, such as TCR-Pred, use molecular structure based on the two-dimensional architecture of CDR3 sequences to predict TCR specificity [36].

Altogether, our exploration of diverse methodologies for inferring TCR specificity combined with a unified TCR database compiling an accurately annotated set of publicly available sequences provides the characteristics of each method that we have summarized in **Table 1**. We believe that our unified TCR database and benchmarking provide a valuable resource for selecting suitable methods for TCR specificity inference, facilitating advancements in understanding the adaptive immune response and developing novel therapies.

**Table 1:** Comparative summary of method characteristics. This table provides a systematic summary of the various characteristics of all the methods evaluated in the study. It categorizes each method’s performance across different metrics into qualitative intervals for ease of interpretation. These intervals are defined as follows: ‘Very Low’ for values in the range of [0-0.2), ‘Low’ for [0.2-0.4), ‘Middle’ for [0.4-0.6), ‘High’ for [0.6-0.8), and ‘Very High’ for [0.8-1]. The last column summarizes the preference of cluster types over the 6 epitopes (**Supplementary Table 5).**

| Methods | Retention | Purity | % clusters with purity >0,9 | Cluster types |
| --- | --- | --- | --- | --- |
| HD | Low | High | Low | Mono/poly-specific |
| LD | Low | Middle | Low | Mono/poly-specific |
| TCRMatch | Very low | **Very high** | High | Mono/poly-specific |
| iSMART | Very low | High | Middle | Mono/poly-specific |
| GIANA | Low | High | Low | Mono/poly-specific |
| clusTCR | Very low | **Very high** | **Very high** | Monospecific |
| GLIPH2 | Low | **Very high** | Middle | Monospecific |
| DeepTCR | **Very high** | High | Low | Polyspecific |
| TCRdist3 | Middle | Middle | Very low | Mono/poly-specific |

## Data availability

The public databases IEDB, McPAS and VDJdb were download between January and March 2023 from the home page of their original web sites. These databases and the final pooled database used in this study are available on GitHub at https://github.com/i3-unit/TCR_Unsupervised_Benchmark. The code used to generate the pooled database, perform the analyses, and create the figures is also accessible via the same repository.

---

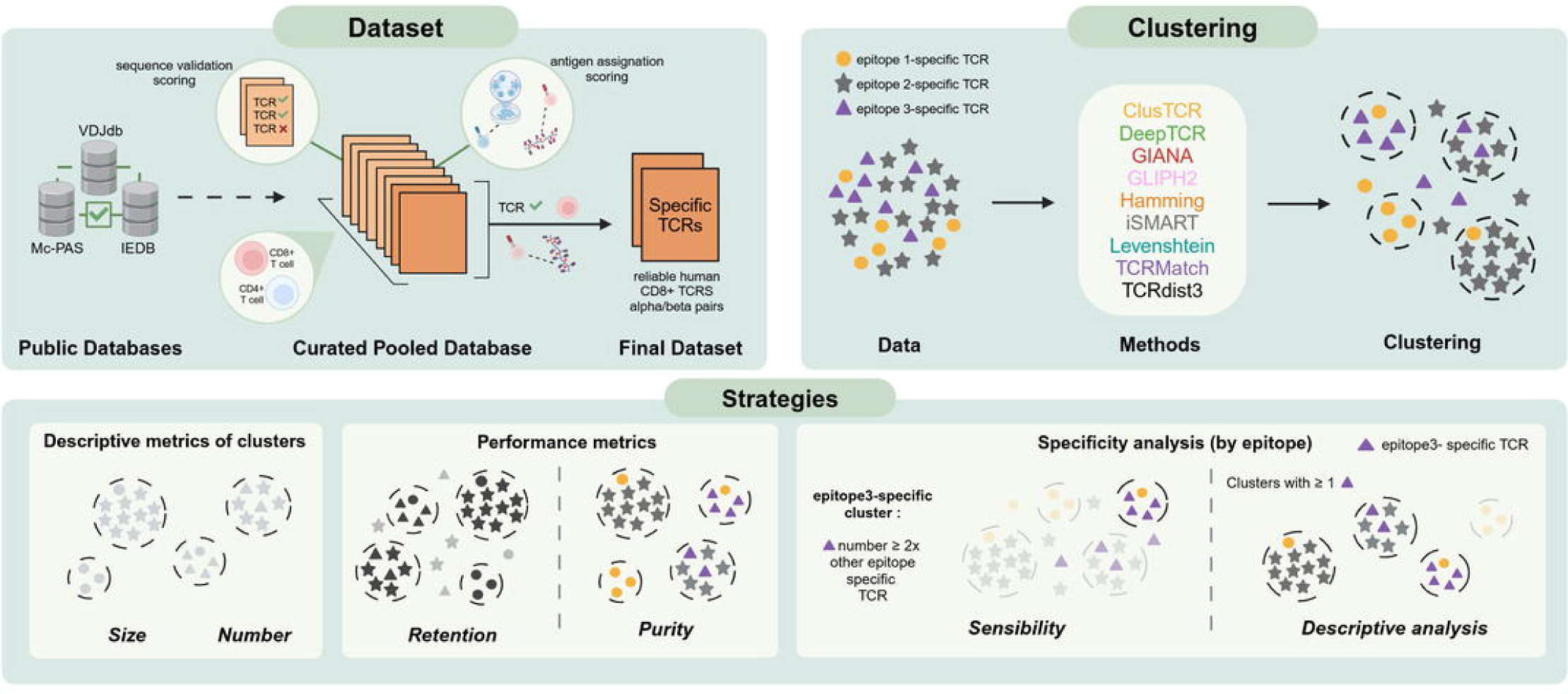

