## Supplemental Figures for "Benchmarking unsupervised methods for inferring TCR specificity"

### *Supplemental Information*

Methods of interest

**- Similarity measures:**

**Levenshtein Distance (LD)** and **Hamming Distance (HD)**: both measure TCR sequence similarity, but LD accommodates sequences of varying lengths through edits (insertions, deletions, substitutions), while HD is limited to sequences of equal length, considering only substitutions. Commonly, sequences differing by one amino acid are deemed similar in both methods (refs).

**TCRMatch** (v.0.1.1). Advances beyond LD and HD by using k-mer approach and the BLOSUM 62 matrix, specifically for TCR β-chain CDR3 (CDR3B) sequences [37]. It identifies and quantifies similarity with a final similarity score ranged from 0 to 1, with 1 indicating a perfect similarity. In our analysis, we set a high similarity threshold above 0.97.

**TCRdist3** (v.0.2.2) [9] is the latest version of TCRdist [38] which offers a comprehensive analysis by including a range of CDR regions and taking into account paired αβ chains, applying a similarity-weighted mismatch distance. It uses the BLOSUM 62 matrix for scoring. For comparing TCRdist3 with the other methods, we applied a hierarchical clustering to its distance matrix. The optimal threshold parameter for the clustering function ('fcluster' from the Python package 'scipy') by maximizing the silhouette score of the clustering solution.

**- Clustering methods:**

**ClusTCR** (v.1.0.2) uses k-means and hashing for speed, connecting similar sequences (HD = 1) within a network, followed by the Markov Clustering Algorithm (MCL) to identify dense substructures representing the clusters [10].

**iSMART** group similar TCRs into antigen-specific clusters through pairwise local alignment using the BLOSUM62 matrix and taking into account the CDR3 sequence length [11]. A depth-first search then identifies connected CDR3 clusters.

**GIANA** (Geometric Isometry based TCR AligNment Algorithm, v.4.1) transforms the CDR3 sequences into numeric vectors using multidimensional scaling (MDS) for fast nearest neighbour searches in the high-dimensional Euclidean space [12]. This method focuses on identifying pre clusters of CDR3 sequences, with additional filtering steps for final cluster formation.

**GLIPH2 is an update to GLIPH** [39] **that** clusters TCRs based on CDR3 similarity, using global and local similarity measures. It analyzes TCR sequences efficiently, overcoming the “small world effect” and employs the Fisher-exact test for cluster confidence [13],  In the column subjet:condition, we have put the following information, “database:CD8+.

**- DeepTCR** (v.2.0) utilizes deep learning for complex TCR sequence analysis. It features a variational autoencoder for feature extraction from CDR3 sequences and offers multiple clustering methods like Phenograph , Hierarchical clustering, and DBSCAN [14]. Hierarchical clustering is specifically used in this study for its efficiency in building nested clusters.

| **Antigen identification method** | **Type of antigen used** | **Antigen identification score** |
|---|---|---|
| Peptide-MHC (pMHC) Xmers sort | NA | 5 |
| In vitro stimulation in CD8 + T-cells | peptide | 4.5 |
| | protein | 4.4 |
| | pathogen | 4.3 |
| | cell line | 4.2 |
| | other | 4.1 |
| In vitro stimulation in CD4+ T-cells | peptide | 3.5 |
| | protein | 3.4 |
| | pathogen | 3.3 |
| | cell line | 3.2 |
| | other | 3.1 |
| Isolated from a specific tissue under a specific pathology / condition | NA | 2 |
| No verified | NA | 1 |
| Missing values | NA | 0 |

**Supplementary Table 1 :** Classification and scoring of antigen identification techniques

| Methods | Input information | Number of clusters (alpha – beta) | Number of small clusters (< 4 sequences) | Number of unique clustered sequences (CDR3a-CDR3b) or pairings |
|---|---|---|---|---|
| **Levenshtein Distance** | CDR3a or CDR3b | 494 (299-195) | 370 (74.8%) | 2248 (1442-806) |
| **Hamming Distance** | CDR3a or CDR3b | 548 (361-187) | 409 (74.6%) | 2564 (1651-913) |
| **TCRMatch** | CDR3a or CDR3b | 285 (188-97) | 253 (88.8%) | 744 (474-270) |
| **iSMART** | CDR3a or CDR3b | 465 (285-180) | 372 (80%) | 1538 (951-587) |
| **GIANA** | CDR3a or CDR3b | 593 (417-175 +1) | 457 (77%) | 2121 (1529-592) |
| **clusTCR** | Paired CDR3a and CDR3b | **111** | 91 (82%) | 426 |
| **GLIPH2** | Paired CDR3a and CDR3b + Vb + Jb + Count + Subject + Condition | 380 | **206 (54.2%)** | 1070 |
| **DeepTCR** | Paired CDR3a and CDR3b + Va + Ja + Vb + Jb + Count | **1628** | 1447 (88.9%) | 4405 |
| **TCRdist3** | Paired CDR3a and CDR3b + Va + Ja + Vb + Jb + Count | 669 | 538 (80.4%) | 2093 |

**Supplementary Table 2 :** Description of required input information and numbers of unique clusters/sequences/pairings formed for each method using the unified database. Input information, number of clusters and number of small clusters (less than 4 sequences) are shown. To note, the numbers of alpha and beta clusters are displayed for methods which do not take into account pairings. Only GIANA has a heterogeneous cluster. Also, the number of unique clustered sequences and pairings are shown. Number of clustered CDR3a and CDR3b is shown for methods which do not consider pairings.

| Methods | Nb of clusters (>3) | Nb of clusters (>5) | Nb of clusters (>10) |
|---|---|---|---|
| Hamming Distance | 139 | 74 | 28 |
| Levenshtein Distance | 124 | 67 | 38 |
| TCRMatch | 32 | 12 | 4 |
| iSMART | 93 | 43 | 10 |
| GIANA | 136 | 66 | 21 |
| clusTCR | 20 | 14 | 6 |
| GLIPH2 | 174 | 94 | 46 |
| DeepTCR | 181 | 63 | 18 |
| TCRdist3 | 131 | 62 | 22 |

**Supplementary Table 3** : Number of clusters formed for each method when considering clusters with size > 3, 5 or 10 sequences/pairs.

| **Epitope** | **Species** | **Count unique CDR3beta** | **Count unique CDR3alpha** | **Count unique CDR3alpha/CDR3beta** |
| --- | --- | --- | --- | --- |
| GILGFVFTL | Influenza | 456 | 484 | 706 |
| NLVPMVATV | CMV | 349 | 325 | 397 |
| GLCTLVAML | EBV | 170 | 145 | 215 |
| FLCMKALLL | TAA | 136 | 134 | 136 |
| NYNYLYRLF | SARS-CoV2 | 46 | 55 | 56 |
| LLFGYPVYV | HTLV-1 | 7 | 4 | 10 |

**Supplementary Table 4 : Detailed overview of selected epitopes and associated species.** CMV: Cytomegalovirus, EBV: Epstein Barr virus, TAA: Tumor associated antigen, SARS-Cov2: Severe acute respiratory syndrome coronavirus 2, HTLV-1: Human T-cell leukemia virus type I

| Methods | GILGFVFTL | NLVPMVATV | GLCTLVAML | FLCMKALLL | NYNYLYRLF | LLFGYPVYV |
|---|---|---|---|---|---|---|
| **HD** | Mono/poly-specific | Polyspecific | Monospecific | Monospecific | Polyspecific | Mono/poly-specific |
| **LD** | Mono/poly-specific | Mono/poly-specific | Monospecific | Monospecific | Mono/poly-specific | Mono/poly-specific |
| **TCRMatch** | Monospecific | Mono/poly-specific | Monospecific | Monospecific | Monospecific | Polyspecific |
| **iSMART** | Mono/poly-specific | Polyspecific | Monospecific | Monospecific | Polyspecific | Mono/poly-specific |
| **GIANA** | Mono/poly-specific | Polyspecific | Monospecific | Monospecific | Polyspecific | Mono/poly-specific |
| **clusTCR** | Monospecific | Monospecific | Monospecific | **X** | Monospecific | Monospecific |
| **GLIPH2** | Monospecific | Monospecific | Monospecific | Monospecific | Monospecific | Monospecific |
| **DeepTCR** | Polyspecific | Polyspecific | Polyspecific | Polyspecific | Polyspecific | Polyspecific |
| **TCRdist3** | Polyspecific | Polyspecific | Polyspecific | Monospecific | Monospecific | Mono/poly-specific |

**Supplementary Table 5 :** Cluster types for the clustering of the 6 epitopes for each method. Barplots representing the percentage of clusters according to the number of specificities for NLVPMVATV, GLCTLVAML, NYNYLYRLF and LLFGYPVYV are not shown in this paper. There are no FLCMKALLL-specific clusters for clusTCR.

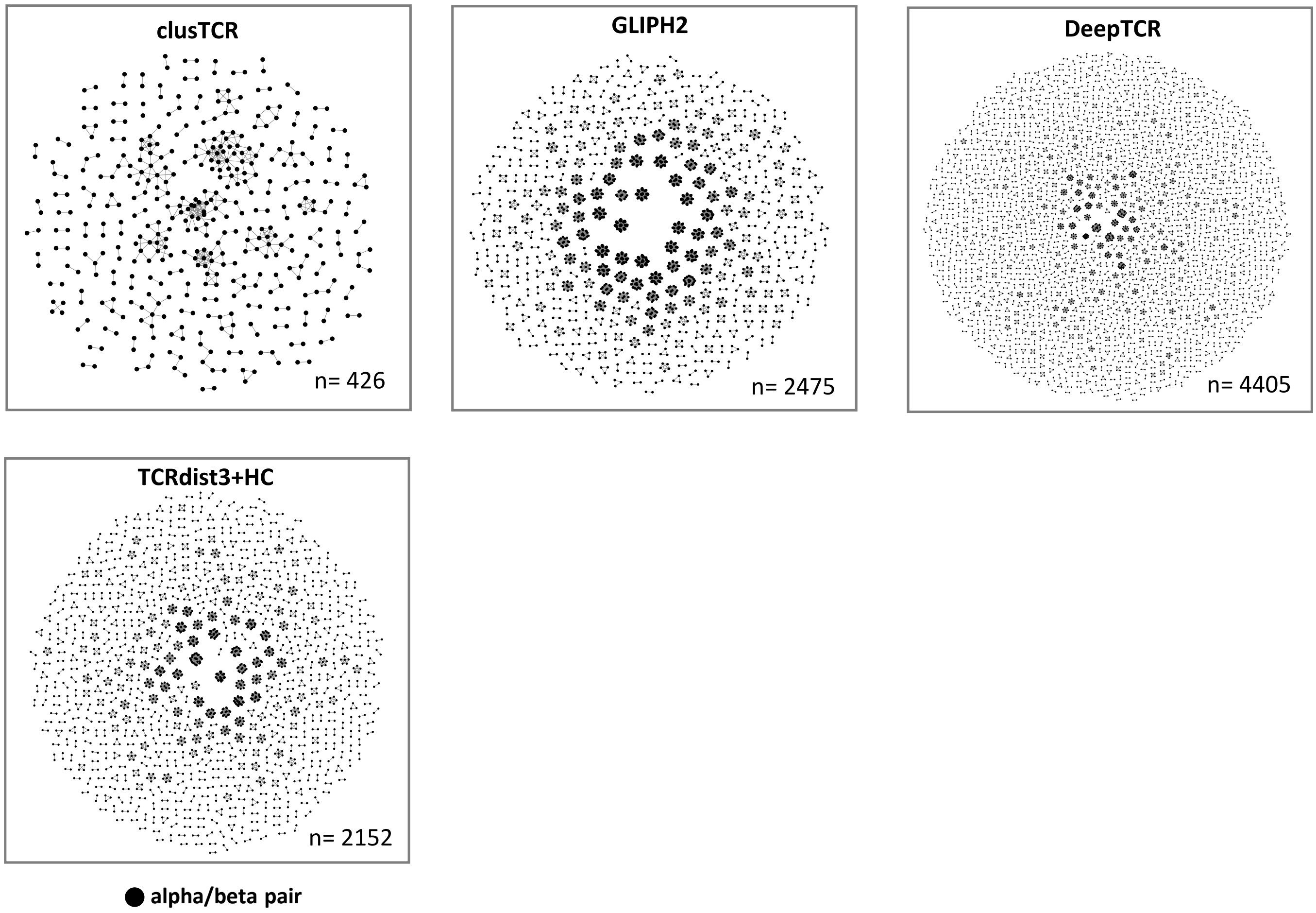


**Supplementary Figure 1:** Networks representing the output clusters for clusTCR, GLIPH2, DeepTCR and TCRdist3. A subnetwork is a cluster. A dot is an alpha/beta pair and it is colored in black. Network for clusTCR: two sequences are linked according the edgelist provided by the tool (HD=1). Network for GLIPH2: all sequences belonging to a cluster are manually linked together. Network for DeepTCR: all sequences belonging to a cluster are manually linked together. Nework for TCRdist3: a hierarchical clustering was performed on the TCRdist3 matrix and two sequences are manually linked together when belonging to the same cluster.

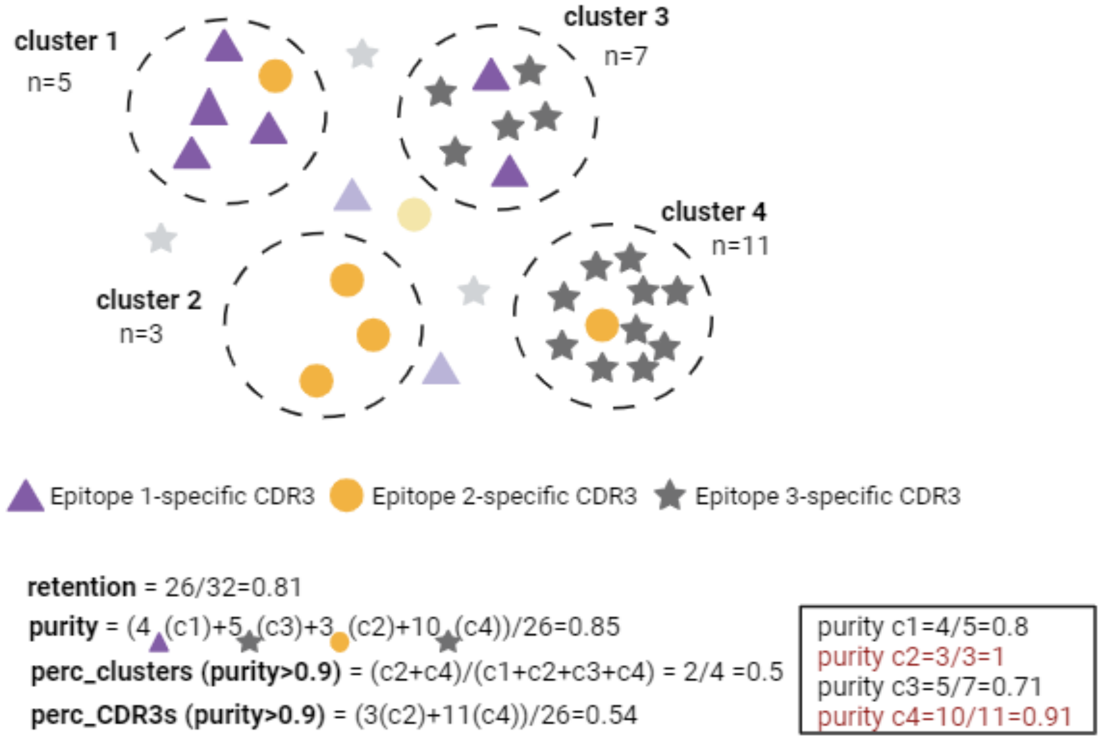


**Supplementary Figure 2 :** Methodology for metric calculation: Schema illustrating the process of calculating clustering evaluation metrics. It provides an example of possible clustering of CDR3 sequences known to bind three distinct epitopes (represented as stars, circles and triangles). Metrics have the following outcomes. Retention: this is the fraction of clustered sequences. In the given example, with 6 sequences outside any clusters, retention is 26/32=0.81. Purity: it assesses the fraction of CDR3s within a single cluster targeting the same epitope. Considering the largest epitope in each cluster, the sum divided by the total number of clustered sequences (4+5+3+10)/26 equals 0.85. Percentage of clusters with a purity over 0.9, individual purity is calculated for each cluster, but only those with a purity over 0.9 are considered. In this example, with individual purities of 0.8 (c1), 1 (c2), 0.71 (c3), and 0.91 (c4), only the c2 and c4 clusters are considered, resulting in 0.5. Percentage of sequences in the above-mentioned high-purity clusters yielding here 3 (c2) + 10 (c2) / 26 = 0.54.

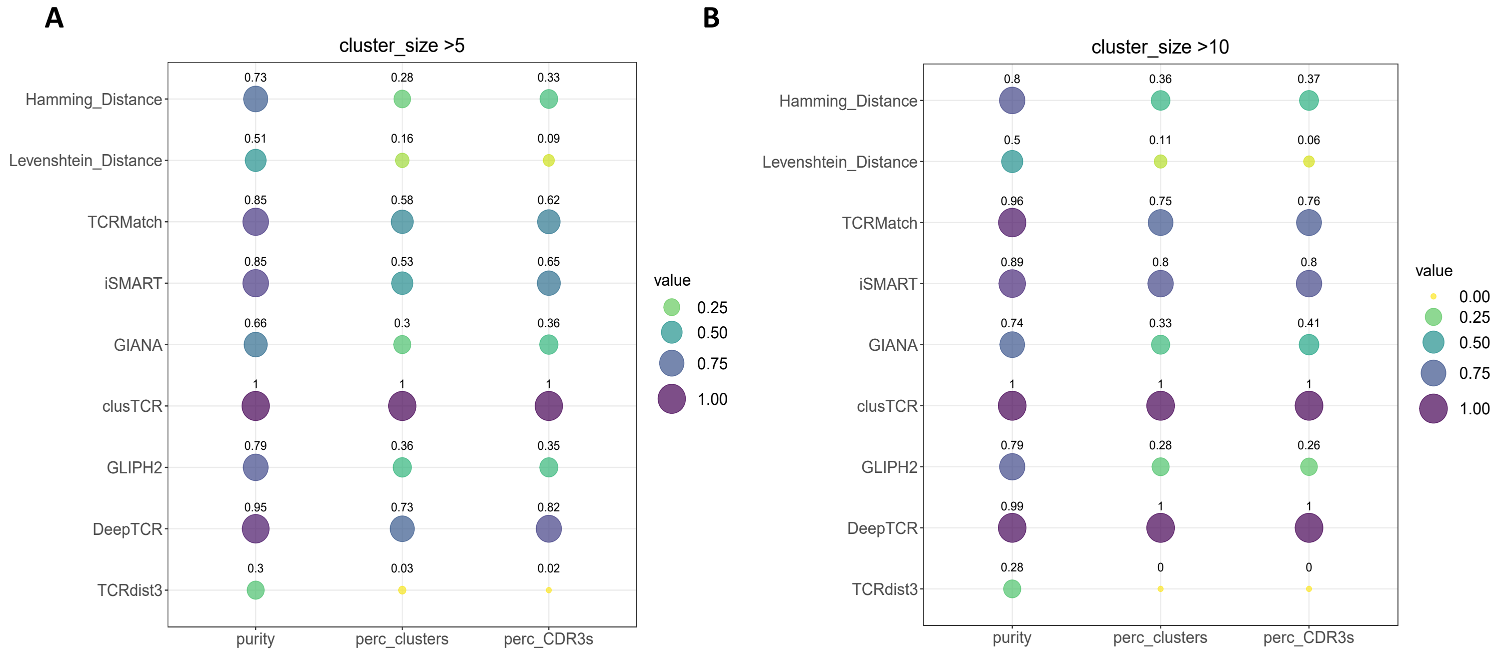


**Supplementary Figure 3 :** Detailed performance of methods. Bubble plots showing the performance evaluation metrics of the nine methods, as in the Figure 3.2.2, with a focus on varying cluster size thresholds: more than 5 and 10 sequences/pairs (A and B respectively).

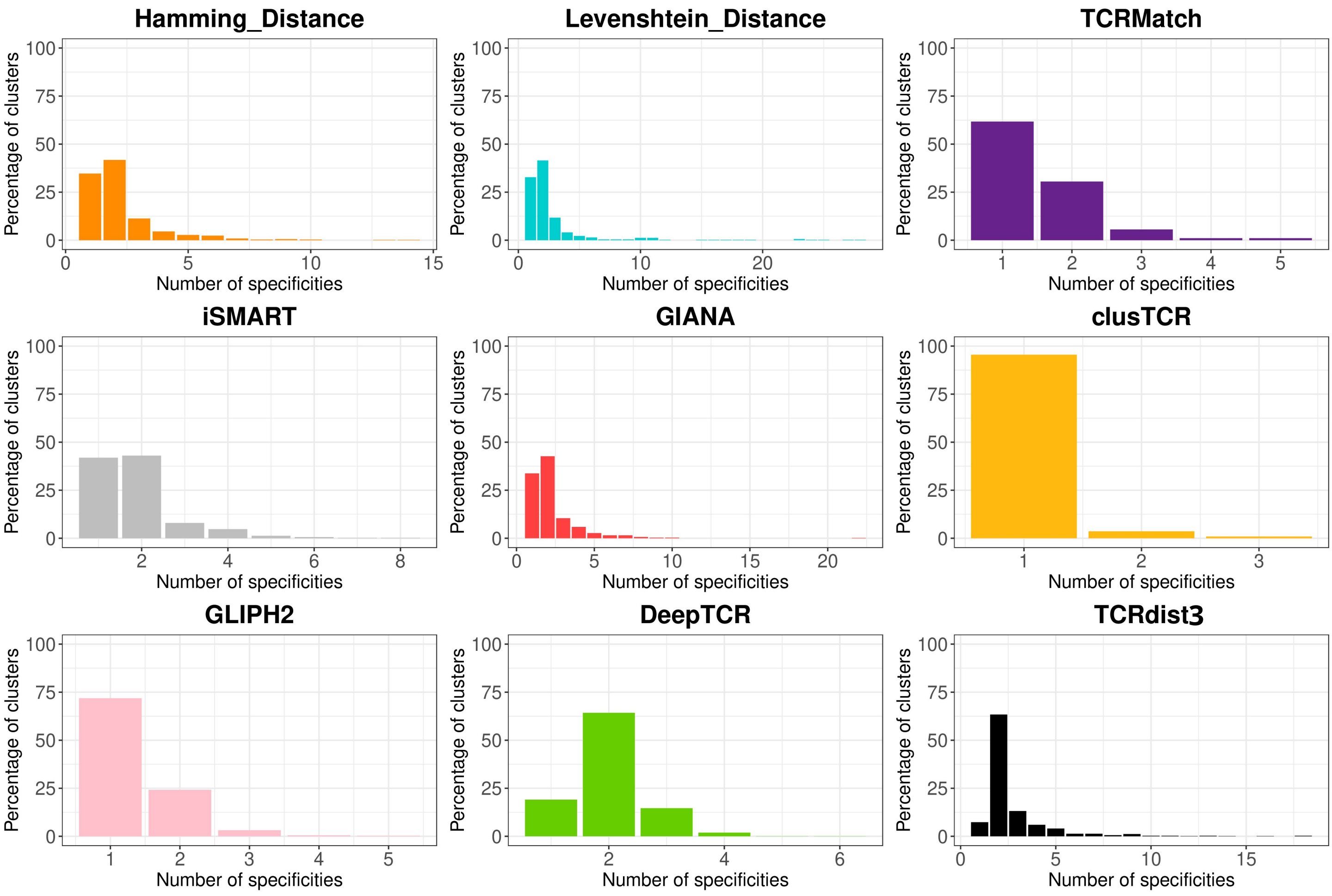


**Supplementary Figure 4 :** Cluster specificity distribution across methods. Percentage of clusters relative to the number of specificities contained within each cluster for each method.

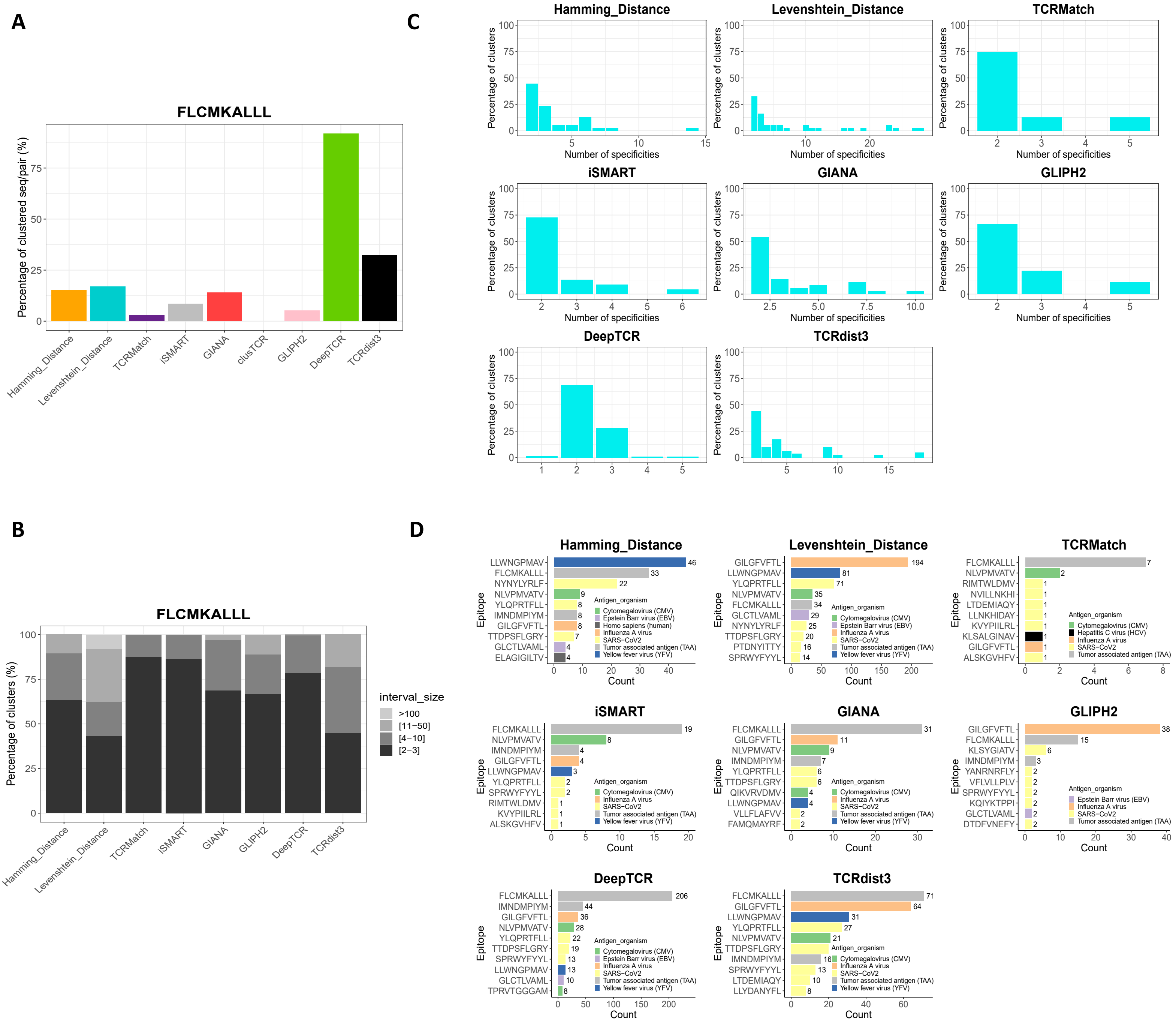


**Supplementary Figure 5 :** In-depth analysis of FLCMKALLL-specific-sequence/pair clustering. (A) Method-dependant clustering efficiency, displaying the percentage of FLCMKALLL-specific-sequences/pairs successfully clustered by each method. (B) FLCMKALLL-specific cluster size distribution for each method. Clusters with at least one specificity GILGFVFTL are considered as specific. (C) Specificity variation in clusters illustrating by the percentage of FLCMKALLL-specific-clusters relative to the number of specificities for each method. (D) Diversity of co-occurring epitopes, illustrated by the top 10 (or less) additional epitopes found within FLCMKALLL-specific clusters, for each clustering method. The epitopes are color-coded according to the antigen organism.

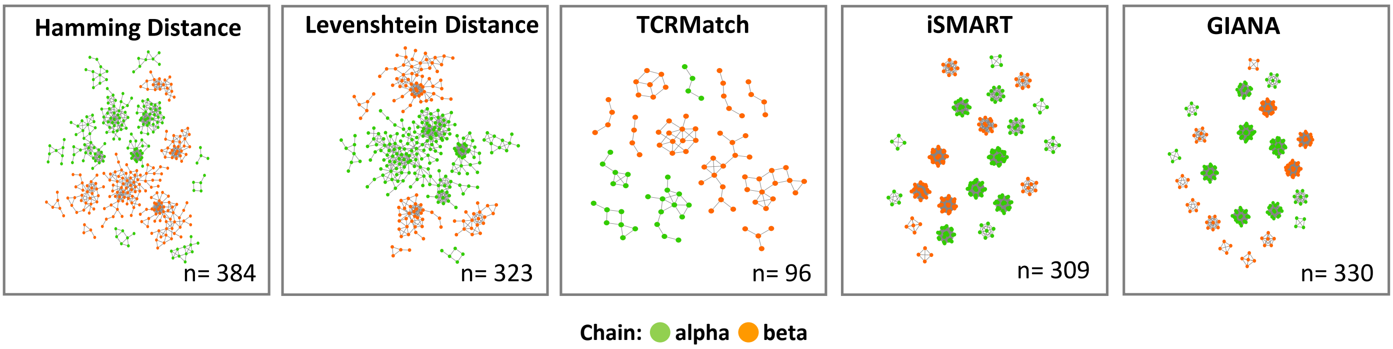


**Supplementary Figure 6:** Visualization of the most specific GILGFVFTL -specific clusters. Network representations of the purest GILGFVFTL-specific clusters (where GILGFVFTL is the major component, being at least twice as prevalent as the second most common epitope withi n the cluster), identified by the first five methods. Each network is color-coded by chain type (CDR3a in green CDR3b in orange).
